# A proof of concept for neutralizing antibody-guided vaccine design against SARS-CoV-2

**DOI:** 10.1101/2020.09.23.309294

**Authors:** Li Zhang, Lei Cao, Xing-Su Gao, Bin-Yang Zheng, Yong-Qiang Deng, Jing-Xin Li, Rui Feng, Qian Bian, Xi-Ling Guo, Nan Wang, Hong-Ying Qiu, Lei Wang, Zhen Cui, Qing Ye, Geng Chen, Kui-Kui Lu, Yin Chen, Yu-Tao Chen, Hong-Xing Pan, Bao-Li Zhu, Cheng-Feng Qin, Xiangxi Wang, Feng-Cai Zhu

## Abstract

Mutations and transient conformational movements of receptor binding domain (RBD) that make neutralizing epitopes momentarily unavailable, present immune escape routes to SARS-CoV-2. To mitigate viral escape, we developed a cocktail of neutralizing antibodies (NAbs) targeting epitopes located on different domains of spike (S) protein. Screening of a library of monoclonal antibodies generated from peripheral blood mononuclear cells of COVID-19 convalescent patients yielded potent NAbs, targeting N-terminal domain (NTD) and RBD domain of S, effective at *nM* concentrations. Remarkably, combination of RBD-targeting NAbs and NTD-binding NAb, FC05, dramatically enhanced the neutralization potency in cell-based assays and animal model. Results of competitive SPR assays and cryo-EM structures of Fabs bound to S unveil determinants of immunogenicity. Combinations of immunogens, identified in NTD and RBD of S, when immunized in rabbits elicited potent protective immune responses against SARS-CoV-2. These results provide a proof-of-concept for neutralization-based immunogen design targeting SARS-CoV-2 NTD and RBD.

**One sentence summary:** Immunogens identified in the NTD and RBD of the SARS-CoV-2 spike protein using a cocktail of non-competing NAbs when injected in rabbits elicited a potent protective immune response against SARS-CoV-2.

## Main Text

Coronavirus disease 2019 (COVID-19), caused by the severe acute respiratory syndrome coronavirus 2 (SARS-CoV-2), continues to spread across the world since December 2019 (*1, 2*). The July 8, 2020 World Health Organization (WHO) Situation Report cited over 11 million COVID-19 cases and 539,000 deaths. These numbers continue to rise daily (*3*). Concerningly, a variant of the SARS-CoV-2 carrying D614G spike mutation, which seemingly enhances the infectivity has been documented and is fast becoming the dominant strain of SARS-CoV-2 globally (*4*). Safe and effective preventive as well as therapeutic measures are urgently needed to bring the ongoing pandemic of COVID-19 under control (*5*). Over the past two months, experimental strategies based on eliciting neutralizing antibodies (NAbs) *via* immunization of potential vaccine candidates and passive administration of NAbs have shown promise in protecting and curing SARS-CoV-2-challenged nonhuman primates (*6-8*). The successes of these studies highlight the importance of screening and identification of immunogens capable of eliciting high NAb titers. Furthermore, NAbs elicited by immunogens differ significantly in their abilities to neutralize SARS-CoV-2 and conferring protection. Therefore, a deep understanding of the nature of NAbs capable of potently neutralizing SARS-CoV-2 and their epitopes could guide new approaches for the development of vaccines.

The coronavirus spike (S) protein is a multifunctional molecular machine that facilitates viral entry into target cells by engaging with cellular receptors and determines to a great extent cell tropism and host range (*9*). Coronaviruses S proteins are processed into S1 and S2 subunits by host proteases, among which S1 is responsible for receptor binding, while the S2 subunit mediates membrane fusion (*10*). The S1 subunit typically possesses two types of domains capable of binding to host cell receptors. For instance, some betacoronaviruses use the N-terminal domain (NTD) of their S1 subunit to bind sialic acids located on the glycosylated cell-surface receptor (*11*). Similarly, the betacoronavirus murine hepatitis virus uses its NTD for binding the protein receptor CEACAM1 (*12*). In contrast to this, SARS-CoV and SARS-CoV-2 use the C-terminal domain of their S1 subunit for binding to their protein receptor hACE2 (*13*). Although it remains unknown whether the NTD is involved in the entry of SARS-CoV-2 into host cells, recent studies have revealed that antibodies targeting the NTD exhibit potent neutralizing activities against SARS-CoV-2 and MERS-CoV infections (*14, 15*). Abrogation of the crucial role played by the S in the establishment of an infection is the main goal of therapies based on neutralizing antibodies and the focus of antibody-based drug and vaccine design. More recently, a number of RBD-targeting NAbs against SARS-CoV-2, which block the binding of the S trimer to hACE2 have been reported and characterized (*16-21*). Stochastic conformational movements of the RBD transiently expose or hide the determinants of receptor binding and some key neutralizing epitopes, which might open up fortuitous escape routes for the virus. Furthermore, antibody-mediated selective pressure is known to lead to antigenic drift within the RBD, resulting in the accumulation of mutations that hamper neutralization by antibodies (*22*). To address these issues related to SARS-CoV-2 neutralization, administration of a cocktail of NAbs targeting both the RBD and non-RBD regions, rather than using a single NAb, could potentially increase the potency of protection *via* binding of NAbs to multiple domains of S, thereby preventing escape of the viral particles from the NAbs. In context with this, the immunogenic characteristics of the antigens targeted by potential NAb cocktails and their structural features can inform strategies for the development of vaccines and therapeutics against COVID-19.

One of the prospective goals of this study was to generate a large and diverse collection of human NAbs targeting multiple domains of S so as to allow for the formulation of a cocktail of highly potent antibodies that could simultaneously bind to the various regions of S. For this, we first established antigen-binding fragment (Fab) phage display libraries from peripheral blood mononuclear cells (PBMCs) of 5 COVID-19 convalescent patients. After three rounds of panning, ∼350 randomly picked colonies were screened by enzyme-linked immunosorbent assay (ELISA) for binding to the SARS-CoV-2 S trimer. A set of 202 positive Fab clones exhibiting tight binding to SARS-CoV-2 were selected for sequencing and further analysis (Fig. 1A). We evaluated the ability of these Fabs to bind to recombinant SARS-CoV-2 NTD, RBD or S2 proteins and observed that 40 (∼20%), 117 (58%) and 45 (22%) of these monoclonal antibodies (mAbs) recognized the NTD, RBD and S2, respectively (Fig. 1A and 1B). We then narrowed down our selection of candidates for the development of a cocktail to 10 antibodies picked from each of these three groups based on their binding affinities and genetic diversity assessed from the phylogenetic analysis performed using the amino acid sequences of the VHDJH and VLJL regions (*23*) (Fig. 1A, fig. S1-S2 and Table S1). Among those selected, 3 (named FC01, FC08 and FC11) target the RBD, 3 (named FC05, FC06 and FC07) recognize the NTD and 4 (named FC118, FC120, FC122 and FC124) are S2-specific mAbs (Fig. 1C and 1D). To find out whether these antibodies cross-react with SARS-CoV and MERS-CoV, we firstly assessed the binding capacity of these mAbs to RBDs, NTDs and S2s from SARS-CoV-2, SARS-CoV and MERS-CoV by ELISA (Fig. 1C). FC11 and FC07 showed cross-binding to SARS-CoV RBD and NTD, respectively. Expectedly, the four S2-directed mAbs interacted with S2s from all the 3 viruses, of which FC122 bound weakly to SARS-CoV and MERS-CoV (Fig. 1C). Surface plasmon resonance (SPR) assays demonstrated that all 10 mAbs exhibit tight bindings to SARS-CoV-2 with affinities in the range of 0.3-62 nM (Fig. 1D). Interestingly, binding affinities of S2-targeting mAbs were relatively weaker than those of RBD- or NTD-targeting antibodies (Fig. 1D and fig. S3).

**Figure 1.**
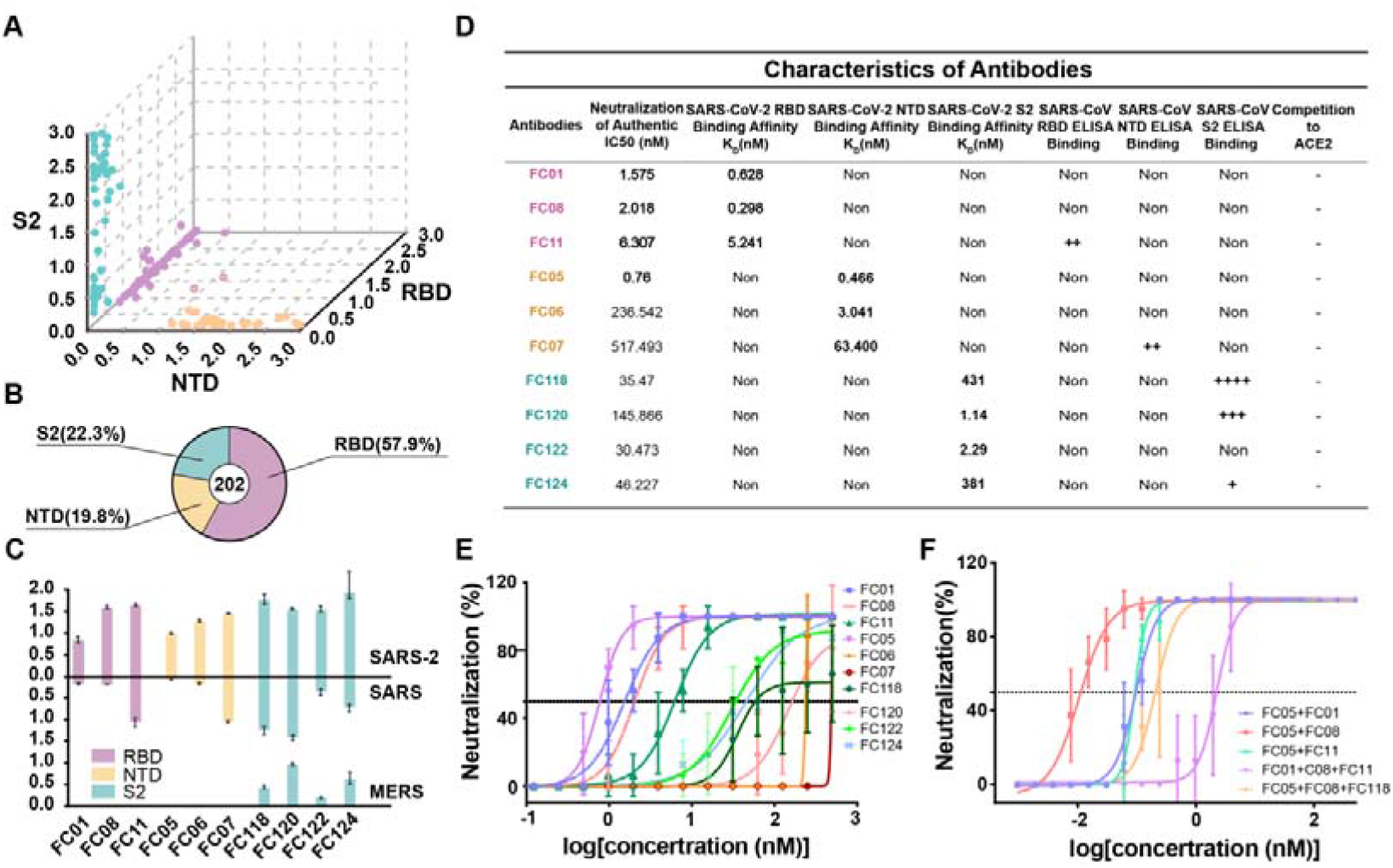
Identification and characterization of SARS-CoV-2 S-targeting neutralizing antibodies. (A) Characteristics of antibodies binding to SARS-CoV-2 RBD, NTD or S2 by ELISA. SARS-CoV-2 RBD-, NTD- and S2-specific mAbs are colored in pink, wheat and pale cyan dots, respectively. A number of mAbs that exhibit non-specific bindings to both SARS-CoV-2 RBD and NTD are presented in wheat dots with pink outlines. (B) Proportion of SARS-CoV-2 S-specific antibodies targeting each of the indicated domains. (C) Bar graph depicting the binding of 10 representative mAbs (FC01, FC05, FC06, FC07, FC08, FC11, FC118, FC120, FC122 and FC124) to S proteins of SARS-CoV-2, SARS-CoV and MERS-CoV using ELISA assays (shown as mean ± S.D. of values derived from experiments conducted in triplicate). (D) Summary of the performance of the representative 10 mAbs in the indicated assays. *In vitro* neutralization activities of 10 individual mAbs (E) or the cocktail of antibodies (F) against SARS-CoV-2 in Vero-E6 cells. Neutralizing activities are represented as mean ± SD. Experiments were performed in triplicates

The effectiveness of the neutralization abilities of the 10 mAbs against SARS-CoV-2 infection when tested using Vero-E6 cells revealed that all 10 showed neutralizing activities with IC50 values ranging from 0.8-520 nM, among which the 3 RBD-targeting and 1 NTD-binding (FC05) mAbs potently neutralized virus at several nM levels (Fig. 1E). These results, together with the results of the binding site studies, allowed us to rationally evaluate the neutralization potency of the NTD-targeting FC05 in combination with the RBD-targeting NAbs. Not surprisingly, the combination of any one of the RBD-targeting NAbs and FC05 enhanced the neutralization potency dramatically when compared to neutralization performed by using individual NAbs under identical conditions (Fig. 1F). Notably, the cocktail consisting of FC05 (NTD-binding) and FC08 (RBD-binding) yielded the strongest neutralizing activity with an IC_50_ value as low as 15 pM, which was better than the cocktail consisting of FC05 and FC01 as well as other combinations of 3 or 4 NAbs (Fig. 1F). Although more recently, synergistic effects between pairs of non-competing RBD-targeting NAbs have been reported for SARS-CoV-2 (*19,22,24*), our cocktail of FC05 and FC08 that bind to different domains of the S trimer provides a proof-of-concept for neutralization-based immunogen design targeting both SARS-CoV-2 NTD and RBD domains.

Next we sought to assess the *in vivo* protection efficacy of these NAbs against a SARS-CoV-2 challenge. A newly established mouse model based on a SARS-CoV-2 mouse adapted strain MASCp6 (*25*) was used to evaluate potential prophylactic and therapeutic efficacy of these NAbs. BALB/c mice were administered a single dose of 20 mg/kg of FC05 or FC08 or a cocktail of FC05 (NTD-binding) and FC08 (RBD-binding) either 12 h before (day −0.5) or 0.5 day (day 0.5) after viral challenge with 2 × 10^4^ PFU of MASCp6 (BetaCoV/Beijing/IMEBJ05-P6/2020) (Fig. 2A). Animals were sacrificed at day 3 for detecting viral loads and examining the pathology of the lungs and tracheas. The number of viral RNA copies estimated in the lungs and tracheas revealed that, in prophylactic settings, a treatment with either individual NAbs or the cocktail led to a 3-4 log reduction of viral loads in both lungs and tracheas at day 3 when compared to the PBS-treated group. Notably, a synergistic protective efficacy was observed for the cocktail (Fig. 2B and 2C). The estimated viral loads from the lungs of groups belonging to therapeutic settings showed similar levels as those observed for the groups of the prophylactic settings, however, the viral loads from the tracheas differed for both the groups. An ∼10-fold higher titer was observed for the groups in therapeutic settings (Fig. 2B and 2C). Histopathological examination revealed a typical interstitial pneumonia, including widening of alveolar septum, vasodilation, hyperemia and edema, accompanied by a large number of monocytes and lymphocytes and a small number of lobulated granulocytes and other inflammatory cell infiltration in mice belonging to the PBS control group (Fig. 2D). In contrast, no obvious lesions of alveolar epithelial cells or focal hemorrhage were observed in the lung sections from either of the antibody-treated groups at day 3 (Fig. 2D).

**Figure 2.**
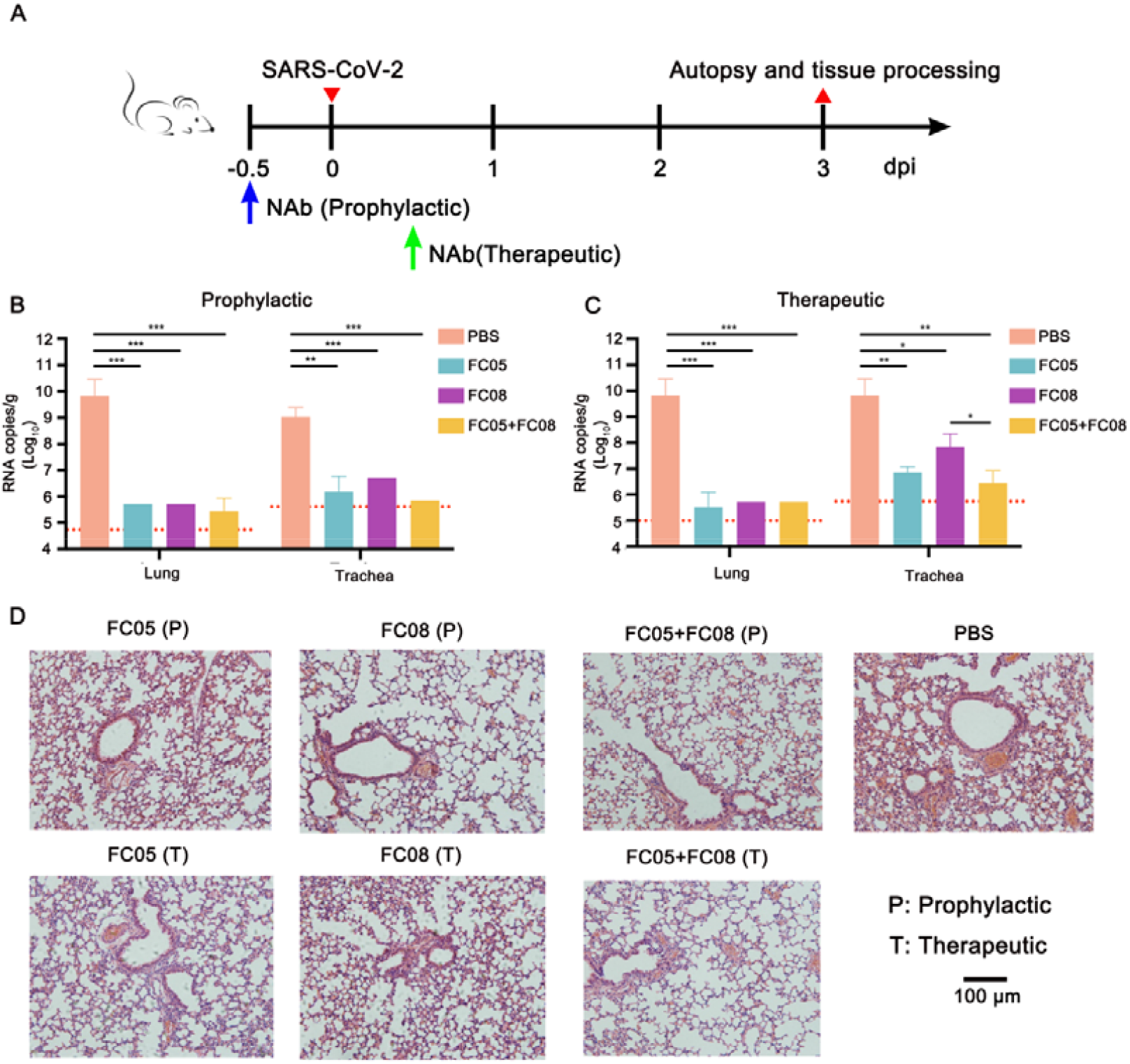
Prophylactic and therapeutic efficacy of FC05 or FC08 or the cocktail of antibodies of FC05 and FC08 in SARS-CoV-2 susceptible mice model. (A) Experimental design for therapeutic and prophylactic evaluations of FC05 or FC08 or the cocktail of antibodies consisting of FC05 and FC08 by using a mouse adapted SARS-CoV-2 virus (MASCp6) in mice model. Group of BALB/c mice were infected intranasally with 2×10^4^ PFU of MASCp6. A dose of 20 mg/kg of antibody was administrated intraperitoneally at 12 hours before infection (the prophylactic group, P) or at 2 hours after infection (the therapeutic group, T). PBS injection was used as a negative control group. Then, the lung and the trachea tissues of mice were collected at 3 and 5 dpi for virus load measurement and histopathological analysis. (B) and (C) Virus loads of lung and trachea tissues at 3 dpi in mouse model. The viral loads of the tissues were determined by qRT-PCR (*P<0.05; **P<0.01; ***P<0.001). Data are represented as mean ± SD. Dashed lines represent the limit of detection. (D) Histopathological analysis of lung and trachea samples at 3 dpi. Scale bar: 100 μm.

To gain a better understanding of the synergy observed during the neutralization of SARS-CoV-2 by a cocktail of NAbs, we performed competitive SPR assays. The results of the assays were expected to reveal whether the NAbs recognize the same or different patches of the epitopes. As expected, the binding of the NTD-specific FC05 does not affect the attachment of any of the three RBD-specific NAbs to the SARS-CoV-2 S trimer, explaining the cooperativity in the antibodies of the cocktail as they bind simultaneously to distinct domains (Fig. 3A). Conversely, the 3 RBD-targeting NAbs competed with each other for binding to the SARS-CoV-2 S trimer (Fig. 3B), which may imply that these RBD-targeting antibodies recognize similar epitopes or their epitopes overlap partially. Lastly, none of the 4 S2-specific NAbs were capable of blocking the interactions between soluble hACE2 and the SARS-CoV-2 S trimer (Fig. 3C). To decipher the nature of the epitopes and the mechanism of neutralization at the atomic level, we determined cryo-EM structures of a prefusion stabilized SARS-CoV-2 S ectodomain trimer in complex with the Fab fragments of the NAbs. Surprisingly, the structural studies revealed that all the three RBD-targeting NAbs are capable of destroying the S trimer into monomers or irregular pieces. A similar perturbation of the S trimer was observed previously in the studies conducted on the CR3022 antibody (*26*). The NTD-binding FC05 however did neither exhibit any such ability of disrupting the S trimer nor did it affect the viral stability (Fig. S4). To gain a deeper understanding of the epitopes targeted by RBD-binding NAbs that disrupt the S trimer, we used a representative antibody, FC08, for performing a competitive SPR-based epitope binding assay. Recently, we mapped the antigenic sites of 3 well characterized RBD-targeting NAbs, H014, HB27 and P17 (*16,20,21*), which bind epitopes located on one side of the RBD, the apical head of the RBD and the receptor binding motif (RBM), respectively. Using these previously characterized antibodies along with FC08 in the assays revealed that only P17 competes with FC08 for binding to the SARS-CoV-2 S trimer. H014 and HB27, as well as the hACE2 can simultaneously bind to SARS-CoV-2 S trimer together with FC08 (Fig. 3D-3F). This observation coupled with an ability of dispersing the S trimer into monomers, suggests that FC08 probably recognizes a cryptic epitope lying towards the interior of the S trimer (Fig. 3D-3F). Cryo-EM characterization of the S-FC05 complex showed full occupancy where one Fab is bound to each NTD of the homotrimeric S (Fig. 3G). 3D classification revealed that the S trimer adopts a 3-fold symmetrical structure with all three RBDs closed albeit without imposing any symmetry. By applying a C3 symmetry, we reconstructed the cryo-EM structure of the complex at an overall resolution of 3.4 Å. A “block-based” reconstruction approach was used to improve the local resolution (3.9 Å) of the map around the binding interface between NTD and FC05 (Fig. 3G, fig. S5-S7 and table S2). Interestingly, the binding mode of FC05 resembles with that of 4A8, a recently reported SARS-CoV-2 NAb (*14*) (fig. S8). Similar to 4A8, FC05 recognizes a conformational epitope formed by elements of the N3 and N5 loops located on the NTD with a buried surface area of ∼700 Å^2^. The essential epitope contains 12 residues, among which all 12 residues (100 %) are not conserved between SARS-CoV and SARS-CoV-2, explaining FC05’s virus-specific binding and neutralization activities (Fig. 3H-3I and fig. S9). The paratope of FC05 is composed of 4 complementarity-determining regions (CDR) loops: CDRL2 (residues 49-55), CDRH1 (residues 29-33), CDRH2 (residues 50-59) and CDRH3 (residues 99-106) (table S3). Extensive hydrophobic and hydrophilic interactions facilitate the tight binding between FC05 and the NTD. Analysis of the structures also provides structural basis for rationalizing the cooperativity observed when both FC05 and FC08 are used for neutralizing SARS-CoV-2.

**Figure 3.**
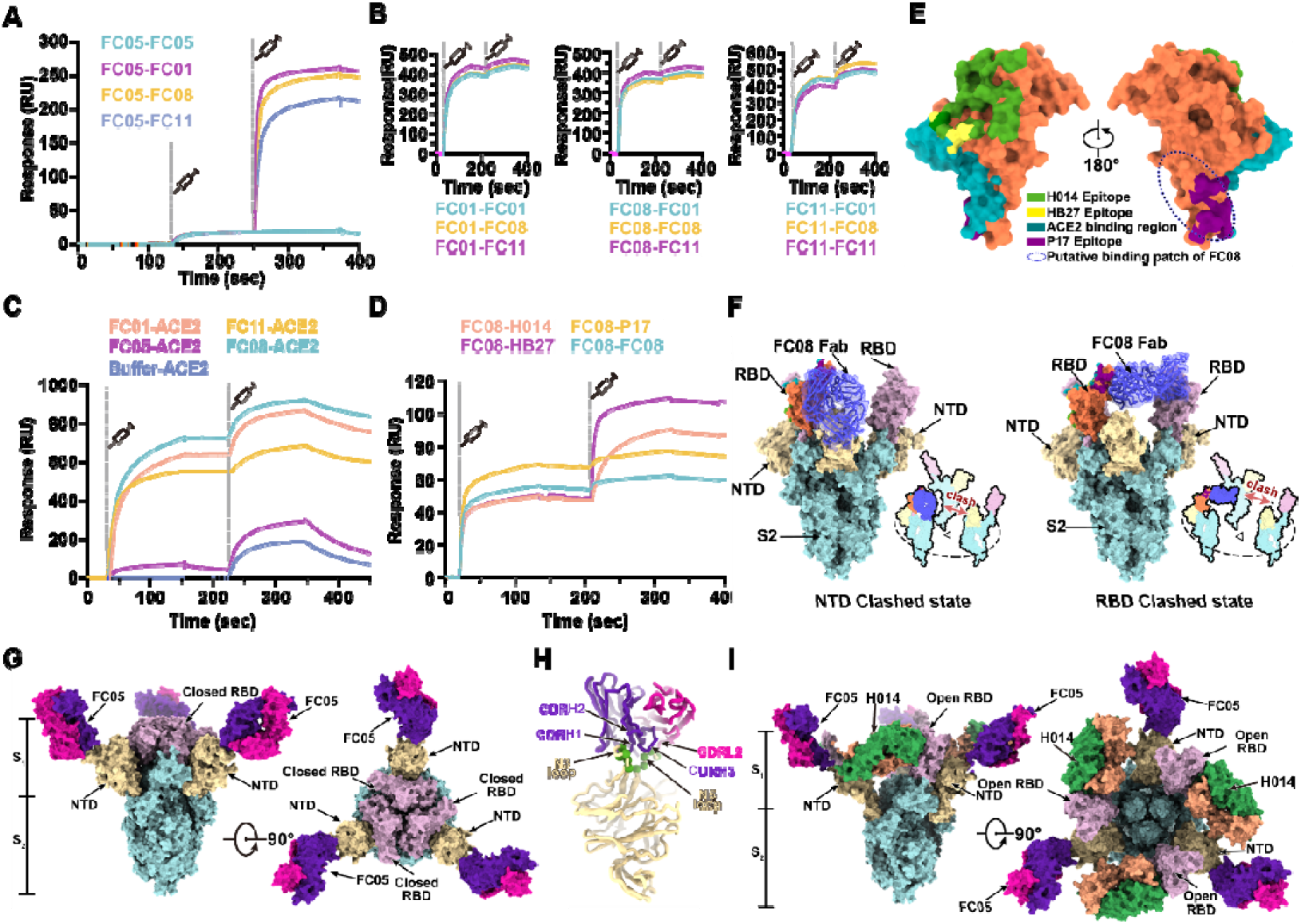
Epitope mapping of SARS-CoV-2 NAbs. (A) SPR kinetics of simultaneous binding of FC05 and 3 RBD-directed NAbs to SARS-CoV-2 S trimer. (B) SPR-based competitive binding of 3 RBD-directed NAbs to SARS-CoV-2 S trimer. SARS-CoV-2 S trimer was initially immobilized onto the sensor. 1 NAb was first injected, followed by the other 2, which indicated these 3 RBD-directed NAbs competed with each other for binding to the SARS-CoV-2 S trimer. (C) Binding of any one RBD-directed NAb blocks the interactions between ACE2 and SARS-CoV-2 S trimer assessed by competitive SPR. (D) Competitive SPR-based epitope mapping of FC08 through 3 recently well characterized RBD-targeting SARS-CoV-2 NAbs, H014, HB27 and P17. The results indicate P17 competes with FC08 for binding to the SARS-CoV-2 S trimer, while H014 and HB27 are capable of simultaneously binding to SARS-CoV-2 S trimer together with FC08. (E) FC08 epitope analysis on the RBD surface. The epitope clusters of H014, P17 and HB27, and the binding region for ACE2 are shown in indicated colors. Putative epitopes are indicated by dashed lines. (F) Two putative models for FC08 binding to SARS-CoV-2 S trimer. Domains of S2 and NTD are colored in cyan and yellow, respectively. The color scheme for the RBD bound FC08 is same as Fig. 3E and the other 2 RBDs are colored in violet. FC08 Fab is presented as blue cartoon with 50% transparent surface. (G) Cryo-EM structure of SARS-CoV-2 S trimer-FC05 complex. Each S monomer is depicted by various colors and the FC05 Fabs are shown in hotpink (light chains) and purpleblue (heavy chains). (H) Interactions between the NTD and FC05. The loops involved in interactions with FC05 are highlighted in green and key CDRs are labeled. (I) Surface representation of the RBD of SARS-CoV-2 S. The areas buried by FC05 is marked by sky blue lines. Sequence identities and differences between the S of SARS-CoV and SARS-CoV-2 are shown in pink and green, respectively, mapped on the surface of SARS-CoV-2 S/RBD.

Most of the potent neutralizing antibodies reported till date target the RBD of CoV (*18-20*). Therefore, a number of RBD-subunit based vaccines for protection against SARS, MERS and COVID-19 are under development (*27-29*). However, RBD-subunit based vaccines could face some critical challenges arising from their relatively low immunogenicity, less diversity within the elicited antibodies and the ensuing potential escape of viral mutants from the antibodies under selective pressure. Our study here, together with other recently published studies, indicates that a subset of NTD-directed antibodies possesses potent neutralizing activities (Fig. 1 and Fig. 2) and that cocktails of antibodies containing NTD-directed as well as RBD-targeting NAbs act in synergy to confer protection against SARS-CoV-2, suggestive of the NTD to be a promising immunogenic partner of the SARS-CoV-2 RBD. To verify this idea, 16 groups of New Zealand rabbits (n=4/group) were injected at day 0, 14 and 28 with various doses of candidate antigen formulations mixed with alum or AS01B adjuvant as follows - 5 μg RBD or 5 μg NTD or 2.5 μg RBD + 2.5 μg NTD; or 20 μg RBD or 20 μg RBD or 10 μg RBD + 10 μg NTD; or 20 μg RBD + 20 μg NTD per dose, 0 μg of antigens in physiological saline as the sham group (Fig. 4). No inflammation or other adverse effects were observed in the animals. Titers of SARS-CoV-2-specific neutralizing antibodies produced by the animals over a period of time (at week 0, 2, 4 and 6) effective in neutralizing the live virus were monitored using microneutralization assays (MN50). Similar to the immune responses elicited by an inactivated SARS-CoV2 vaccine candidate (PiCoVacc) reported by us previously, the neutralizing antibody titer emerged at week 2, surged at week 4 and continued to increase at week 6 (Fig. 4). Perhaps correlated with the less glycosylation and more neutralizing epitopes present in RBD, the RBD induced much higher NAb titers than the NTD at various doses. However, the combination of RBD and NTD exhibited more robust and stable immunogenicity for neutralization compared with a single immunogen consisting of either RBD or NTD at the same dose tested under identical conditions (Fig. 4). Notably, relatively large differences in the ability of individual animals in eliciting NAb titers were observed within the RBD vaccinated groups. Addition of NTD to RBD not only substantially enhanced the NAb titer, but also remarkably decreased the fluctuation in eliciting NAb titer from immunized animals (Fig. 4). Compared to AS01B, alum based adjuvant facilitates the antigens in boosting immune responses at 5 or 20 μg/dose. Administration of higher doses of antigens during immunization (20 μg RBD + 20 μg NTD) with AS01B led to the highest NAb titer, up to ∼1000. In contrast, immunization with higher doses of antigens in conjunction with alum yielded a decreased NAb titer (∼200) compared to the median dose (10 μg RBD + 10 μg NTD), indicative of the need for proper collocation of the adjuvant and various doses of antigen during immunization (Fig. 4).

**Figure 4.**
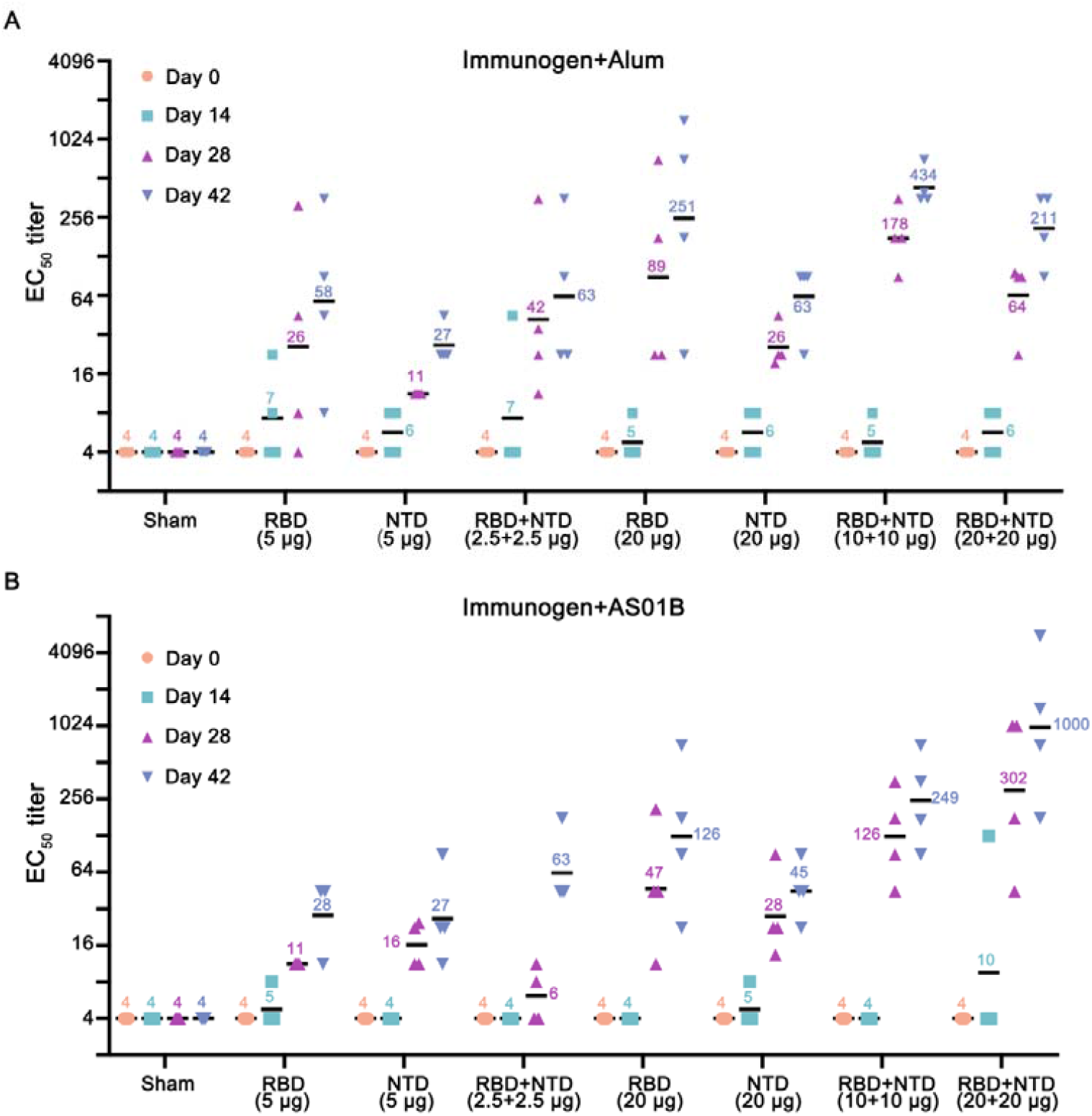
Immunogenic evaluation of candidate antigen formulations mixed with alum or AS01B adjuvant in rabbits. (A-B) Rabbits were immunized intramuscularly with various doses of individual (RBD or NTD) or combined (RBD+NTD) immunogens mixed with alum or AS01B adjuvant or adjuvant only (sham) (n=4). Neutralizing antibody titer against live SARS-CoV-2 was measured. Data points represent mean +/- SEM of individual rabbits from four independent experiments; error bars reflect SEM; horizontal lines indicate the geometric mean titer (GMT) of EC_50_ for each group.

COVID-19 vaccine development is moving at unprecedented speed, with more than 250 candidates under development worldwide. Except for a few inactivated SARS-CoV-2 virus vaccines, most of the candidate vaccines are aimed at the RBD or the spike of SARS-CoV-2 as target immunogen. Selection of the target immunogen is critical for the success of a vaccine, since eliciting a large amount of antibody that binds, but does not neutralize, may lead to low protection or even immunopathology (*30*). The ideal immunogen should elicit high-quality, functionally neutralizing antibodies while avoiding induction of non-neutralizing antibodies. We are now entering a new era of precision vaccinology in which multi-disciplinary techniques provide avenues to rapidly isolate and characterize human NAbs, to define the structural basis of antigenicity, to understand mechanisms of viral neutralization and to guide rational immunogen design. The immunogenic data derived from immunization with a combination of SARS-CoV-2 RBD and NTD here demonstrates the feasibility of eliciting robust targeted immune profiles by using antibody-guided vaccine design and advance us a step forward towards a future of precision vaccines.

## Acknowledgments

We thank Prof. Zihe Rao for valuable discussion on this project and thank Drs. Chunyun Sun, and Changfa Fan for providing critical reagents and thank Drs. Xiaojun Huang, Boling Zhu and Gang Ji for cryo-EM data collection, the Center for Biological imaging (CBI) in Institute of Biophysics for EM work. This work was supported by the National Key Research and Development Program (2020YFA0707500, 2018YFA0900801), the Strategic Priority Research Program (XDB29010000), National Natural Science Foundation of China (NSFC) (grants 82041005, 31900873). Feng-Cai Zhu and Li Zhang were supported by the Research and development project of Jiangsu Province (BE20200601) and The Social Development Project of Jiangsu Province (BE2020720). Xiangxi Wang was supported by Ten Thousand Talent Program and the NSFS Innovative Research Group (No. 81921005). Cheng-Feng Qin was supported by the National Science Fund for Distinguished Young Scholar (No. 81925025) and the Innovative Research Group (No. 81621005) from the NSFC, and the Innovation Fund for Medical Sciences (No.2019-I2M-5-049) from the Chinese Academy of Medical Sciences.

## Author contribution

F-C.Z., X.W. and C-F.Q conceived, designed and supervised the study and wrote the paper. L.Z., X-S.G., B-Y.Z., contributed in construction of the antibody libraries, panning and sequencing of mAbs, generation of mAbs, and the cross-reaction of mAbs. J-X.L. contributed in the paper drafting, plotting histogram, and data interpretation. H-X.P., B-L.Z. collected data. X-L.G., Y.C., performed neutralizing activities assay (CPE) in Vero-E6 cells. Q.B., G.C., K-K. L., evaluated safety and immunogenicity of a recombinant two components candidate COVID-19 vaccine in New Zealand rabbits. C.L. purified proteins, prepared cryo-EM grids and collected cryo-EM data. Y-Q.D. and Q.Y. performed live virus and animal assays; N.W., L.W., L.C. and X.W. processed data. L.C. built and refined the structure model. N.W., L.C. and X.W. analyzed the structures. R.F. performed SPR assay. All authors read and approved the contents of the manuscript.

## Data and materials availability

Cryo-EM density maps have been deposited at the Electron Microscopy Data Bank with accession codes EMD-XXXX (SARS-CoV-2 S-FC05) and EMD-YYYY (SARS-CoV-2 S in complex with FC05 and H014) and related atomic models have been deposited in the protein data bank under accession code 6XXX and 6YYY, respectively.

## References and Notes

1. N. Zhu, D. Zhang et al., A Novel Coronavirus from Patients with Pneumonia in China, 2019. The New England journal of medicine 382, 727–733 (2020); published online EpubFeb 20 (10.1056/NEJMoa2001017).

2. N. Chen, M. Zhou et al., Epidemiological and clinical characteristics of 99 cases of 2019 novel coronavirus pneumonia in Wuhan, China: a descriptive study. Lancet (London, England) 395, 507–513 (2020); published online EpubFeb 15 (10.1016/s0140-6736(20)30211-7).

3. W. H. Organization. (July 8, 2020)

4. B. Korber, W. M. Fischer et al., Tracking changes in SARS-CoV-2 Spike: evidence that D614G increases infectivity of the COVID-19 virus. Cell, (2020); published online Epub2020/07/03/ (https://doi.org/10.1016/j.cell.2020.06.043).

5. K. P. O’Callaghan, A. M. Blatz et al., Developing a SARS-CoV-2 Vaccine at Warp Speed. Jama, (2020); published online EpubJul 6 (10.1001/jama.2020.12190).

6. Q. Gao, L. Bao et al., Rapid development of an inactivated vaccine candidate for SARS-CoV-2. Science, (2020); published online EpubMay 6 (10.1126/science.abc1932).

7. J. Yu, L. H. Tostanoski et al., DNA vaccine protection against SARS-CoV-2 in rhesus macaques. Science, (2020); published online EpubMay 20 (10.1126/science.abc6284).

8. R. Shi, C. Shan et al., A human neutralizing antibody targets the receptor-binding site of SARS-CoV-2. Nature, (2020); published online EpubMay 26 (10.1038/s41586-020-2381-y).

9. F. Li, Structure, Function, and Evolution of Coronavirus Spike Proteins. Annual review of virology 3, 237–261 (2016); published online EpubSep 29 (10.1146/annurev-virology-110615-042301).

10. M. Hoffmann, H. Kleine-Weber et al., SARS-CoV-2 Cell Entry Depends on ACE2 and TMPRSS2 and Is Blocked by a Clinically Proven Protease Inhibitor. Cell 181, 271–280 e278 (2020); published online EpubApr 16 (10.1016/j.cell.2020.02.052).

11. X. Huang, W. Dong et al., Human Coronavirus HKU1 Spike Protein Uses O-Acetylated Sialic Acid as an Attachment Receptor Determinant and Employs Hemagglutinin-Esterase Protein as a Receptor-Destroying Enzyme. Journal of virology 89, 7202–7213 (2015); published online EpubJul (10.1128/JVI.00854-15).

12. J. Shang, Y. Wan et al., Structure of mouse coronavirus spike protein complexed with receptor reveals mechanism for viral entry. PLoS pathogens 16, e1008392 (2020); published online EpubMar (10.1371/journal.ppat.1008392).

13. J. Shang, Y. Wan et al., Cell entry mechanisms of SARS-CoV-2. Proceedings of the National Academy of Sciences of the United States of America 117, 11727–11734 (2020); published online EpubMay 26 (10.1073/pnas.2003138117).

14. X. Chi, R. Yan et al., A neutralizing human antibody binds to the N-terminal domain of the Spike protein of SARS-CoV-2. Science, (2020); published online EpubJun 22 (10.1126/science.abc6952).

15. H. Zhou, Y. Chen et al., Structural definition of a neutralization epitope on the N-terminal domain of MERS-CoV spike glycoprotein. Nature communications 10, 3068 (2019); published online EpubJul 11 (10.1038/s41467-019-10897-4).

16. Y.-Q. D. Zhe Lv, Qing Ye, Lei Cao, Chun-Yun Sun, Changfa Fan, Weijin Huang, Shihui Sun, Yao Sun, Ling Zhu, Qi Chen, Nan Wang, Jianhui Nie, Zhen Cui, Dandan Zhu, Neil Shaw, Xiao-Feng Li, Qianqian Li, Liangzhi Xie, Youchun Wang, Zihe Rao, Cheng-Feng Qin, Xiangxi Wang, Structural basis for neutralization of SARS-CoV-2 and SARS-CoV by a potent therapeutic antibody. bioRxiv : the preprint server for biology, (2020).

17. D. Pinto, Y. J. Park et al., Cross-neutralization of SARS-CoV-2 by a human monoclonal SARS-CoV antibody. Nature 583, 290–295 (2020); published online EpubJul (10.1038/s41586-020-2349-y).

18. A. Z. Wec, D. Wrapp et al., Broad neutralization of SARS-related viruses by human monoclonal antibodies. Science, (2020); published online EpubJun 15 (10.1126/science.abc7424).

19. P. J. M. Brouwer, T. G. Caniels et al., Potent neutralizing antibodies from COVID-19 patients define multiple targets of vulnerability. Science, (2020); published online EpubJun 15 (10.1126/science.abc5902).

20. Y.-Q. D. Ling Zhu, Rong-Rong Zhang, Zhen Cui, Chun-Yun Sun, Chang-Fa Fan, Liangzhi Xie, Youchun Wang, Xiangxi Wang, Cheng-Feng Qin, Double Lock of a Potent Human Therapeutic Monoclonal Antibody against SARS-CoV-2. bioRxiv : the preprint server for biology, (2020).

21. Y. S. Hangping Yao, Yong-Qiang Deng, Nan Wang, Xiao-Feng Li, Yongcong Tan, Na-na Zhang, Luanjuan Li, Cheng-Feng Qin, Xiangxi Wang, Rational Development of a Human Antibody Cocktail that Deploys Multiple Functions to Confer Pan-SARS-CoVs Protection. bioRxiv : the preprint server for biology, (2020).

22. A. Baum, B. O. Fulton et al., Antibody cocktail to SARS-CoV-2 spike protein prevents rapid mutational escape seen with individual antibodies. Science, (2020); published online EpubJun 15 (10.1126/science.abd0831).

23. S. Kumar, G. Stecher et al., MEGA7: Molecular Evolutionary Genetics Analysis Version 7.0 for Bigger Datasets. Molecular biology and evolution 33, 1870–1874 (2016); published online EpubJul (10.1093/molbev/msw054).

24. Y. Wu, F. Wang et al., A noncompeting pair of human neutralizing antibodies block COVID-19 virus binding to its receptor ACE2. Science 368, 1274–1278 (2020); published online EpubJun 12 (10.1126/science.abc2241).

25. H. Gu, Q. Chen et al., Rapid adaptation of SARS-CoV-2 in BALB/c mice: Novel mouse model for vaccine efficacy. bioRxiv, 2020.2005.2002.073411 (2020)10.1101/2020.05.02.073411).

26. J. Huo, Y. Zhao et al., Neutralization of SARS-CoV-2 by Destruction of the Prefusion Spike. Cell host & microbe, (2020); published online EpubJun 19 (10.1016/j.chom.2020.06.010).

27. L. Dai, T. Zheng et al., A Universal Design of Betacoronavirus Vaccines against COVID-19, MERS, and SARS. Cell, (2020); published online EpubJun 28 (10.1016/j.cell.2020.06.035).

28. N. Wang, J. Shang et al., Subunit Vaccines Against Emerging Pathogenic Human Coronaviruses. Frontiers in microbiology 11, 298 (2020)10.3389/fmicb.2020.00298).

29. J. Yang, W. Wang et al., A vaccine targeting the RBD of the S protein of SARS-CoV-2 induces protective immunity. Nature, (2020); published online EpubJul 29 (10.1038/s41586-020-2599-8).

30. B. S. Graham, Rapid COVID-19 vaccine development. Science (New York, N.Y.) 368, 945–946 (2020); published online EpubMay 29 (10.1126/science.abb8923).

31. C. F. Barbas, 3rd, A. S. Kang et al., Assembly of combinatorial antibody libraries on phage surfaces: the gene III site. Proceedings of the National Academy of Sciences of the United States of America 88, 7978–7982 (1991); published online EpubSep 15 (10.1073/pnas.88.18.7978).

32. Z. Chen, X. Ren et al., An elaborate landscape of the human antibody repertoire against enterovirus 71 infection is revealed by phage display screening and deep sequencing. mAbs 9, 342–349 (2017); published online EpubFeb/Mar (10.1080/19420862.2016.1267086).

33. D. N. Mastronarde, Automated electron microscope tomography using robust prediction of specimen movements. Journal of structural biology 152, 36–51 (2005); published online EpubOct (10.1016/j.jsb.2005.07.007).

34. K. Zhang, Gctf: Real-time CTF determination and correction. Journal of structural biology 193, 1–12 (2016); published online EpubJan (10.1016/j.jsb.2015.11.003).

35. S. H. Scheres, Processing of Structurally Heterogeneous Cryo-EM Data in RELION. Methods in enzymology 579, 125–157 (2016)10.1016/bs.mie.2016.04.012).

36. S. H. Scheres, S. Chen, Prevention of overfitting in cryo-EM structure determination. Nature methods 9, 853–854 (2012).

37. Y. Yang, P. Yang et al., Architecture of the herpesvirus genome-packaging complex and implications for DNA translocation. Protein & cell 11, 339–351 (2020); published online EpubMay (10.1007/s13238-020-00710-0).

38. N. Wang, D. Zhao et al., Architecture of African swine fever virus and implications for viral assembly. Science 366, 640–644 (2019); published online EpubNov 1 (10.1126/science.aaz1439).

39. N. Wang, W. Chen et al., Structures of the portal vertex reveal essential protein-protein interactions for Herpesvirus assembly and maturation. Protein & cell 11, 366–373 (2020); published online EpubMay (10.1007/s13238-020-00711-z).

40. A. Kucukelbir, F. J. Sigworth et al., Quantifying the local resolution of cryo-EM density maps. Nature methods 11, 63–65 (2014).

41. L. A. Kelley, S. Mezulis et al., The Phyre2 web portal for protein modeling, prediction and analysis. Nature Protocols 10, 845–858 (2015); published online EpubJun (10.1038/nprot.2015.053).

42. E. F. Pettersen, T. D. Goddard et al., UCSF Chimera—a visualization system for exploratory research and analysis. Journal of computational chemistry 25, 1605–1612 (2004).

43. A. Brown, F. Long et al., Tools for macromolecular model building and refinement into electron cryo-microscopy reconstructions. Acta Crystallographica Section D-Structural Biology 71, 136–153 (2015); published online EpubJan (10.1107/S1399004714021683).

44. P. V. Afonine, R. W. Grosse-Kunstleve et al., Towards automated crystallographic structure refinement with phenix. refine. Acta Crystallographica Section D: Biological Crystallography 68, 352–367 (2012).

45. V. B. Chen, W. B. Arendall et al., MolProbity: all-atom structure validation for macromolecular crystallography. Acta Crystallographica Section D: Biological Crystallography 66, 12–21 (2010).

46. P. Gouet, E. Courcelle et al., ESPript: analysis of multiple sequence alignments in PostScript. Bioinformatics 15, 305–308 (1999).

